# Experimental Context Shapes PRR-Mediated Immune Output Sensitivity in *Arabidopsis*

**DOI:** 10.64898/2026.04.29.721445

**Authors:** Alba Moreno-Pérez, Hanxu Sha, Gitta Coaker

**Author notes:** **Material distribution footnote:** The author responsible for distribution of materials integral to the findings presented in this article in accordance with the policy described in the Instructions for Authors (https://academic.oup.com/plcell/pages/General-Instructions) is: Gitta Coaker.

## Abstract

Pattern recognition receptors (PRRs) mediate plant immune responses by detecting extracellular immunogenic patterns, including microbe-associated molecular patterns (MAMPs). PRR signaling is commonly assessed using assays such as reactive oxygen species (ROS) bursts, cytosolic calcium influx, mitogen-activated protein kinase (MAPK) activation, and seedling growth inhibition (SGI), which are performed in distinct experimental systems, including seedlings grown on artificial media and soil-grown rosettes. Here, we systematically compare receptor kinase immune outputs triggered by the bacterial MAMPs elf18 and flg22 in *Arabidopsis thaliana* seedlings and rosettes across a range of concentrations. Rosettes exhibited greater sensitivity than seedlings in ROS assays, whereas cytosolic calcium responses measured using the Aeq^cyt^/pMAQ2 reporter were stronger in seedlings, correlating with reduced reporter transcript accumulation in rosette tissue. MAPK activation was consistently stronger in rosettes, whereas SGI assays revealed higher sensitivity to elf18 than flg22 in seedlings despite flg22 inducing stronger early signaling outputs. Together, these results demonstrate that canonical PRR-mediated immune outputs are differentially sensitive to experimental context and should not be interpreted as interchangeable measures of immune activation. These findings highlight the importance of considering experimental conditions when comparing immune responses across assays and developmental stages.

## Introduction

All classes of pathogens can infect plants, and losses due to pests and diseases range from 8-41% across major commodities (Savary et al. 2019). To defend against these threats, plants recognize pathogens or pathogen-induced damage using surface-localized pattern recognition receptors (PRRs) with receptor kinase or receptor protein architecture (Zhou and Zhang 2020; Ngou et al. 2024). These PRRs detect microbial features as non-self, known as microbe-associated molecular patterns (MAMPs), and induce defense responses (Tanaka and Heil 2021; Ngou et al. 2022; Li et al. 2024). Some PRRs can also recognize extracellular pathogen effector proteins (Ngou et al. 2022).

PRR activation induces multiple defense responses, including reactive oxygen species (ROS) production, apoplast alkalization, mitogen-activated protein kinase (MAPK) activation, hormone biosynthesis, transcriptional reprogramming, and callose deposition, culminating in disease resistance (Boller and Felix 2009; Yu et al. 2017). Sustained MAMP exposure can also reduce plant growth, commonly assessed using seedling growth inhibition (SGI) in liquid media. These responses differ in their spatial and temporal dynamics, occurring within seconds to minutes or over hours to days (Boller and Felix 2009; Yu et al. 2017).

Calcium acts as a key second messenger in plant immunity and is rapidly induced following PRR activation through ion fluxes across the plasma membrane (Ghosh et al. 2022; Wdowiak et al. 2024). In parallel, PRR signaling triggers an apoplastic ROS burst, mediated by NADPH oxidases such as RBOHD in *Arabidopsis thaliana* (hereafter referred to as *Arabidopsis*), leading to hydrogen peroxide accumulation (Petrov and Van Breusegem 2012; Mittler et al. 2022). Calcium and ROS signaling are tightly interconnected; calcium-dependent activation of RBOHD promotes ROS accumulation, while ROS can reinforce calcium signaling and contributes to local and systemic defense responses (Dubiella et al. 2013; Kadota et al. 2015; Evans et al. 2016; Choi et al. 2017).

Activation of MAPK cascades is another conserved immune response downstream of PRR signaling (Meng and Zhang 2013; Majeed et al. 2022; Sun and Zhang 2022). MAPKs transduce MAMP perception signals and interface with WRKY transcription factors to drive transcriptional reprogramming toward defense (Mao et al. 2011; Sun and Zhang 2022). In *Arabidopsis*, MAMP recognition primarily activates two parallel MAPK cascades, MEKK3/5–MKK4/5–MPK3/6 and MEKK1–MKK1/2–MPK4 (Bi et al. 2018; Sun and Zhang 2022; Yan et al. 2025). The MEKK1–MKK1/2–MPK4 pathway plays a predominant role in immunity by promoting defense gene expression, phytoalexin and ethylene biosynthesis, and stomatal immune responses (Sun and Zhang 2022). MAPK activation occurs within 5-15 minutes after PRR activation (Bethke et al. 2012).

Two of the most extensively studied PRRs are the leucine-rich repeat receptor kinases FLAGELLIN SENSING 2 (FLS2) and ELONGATION FACTOR-Tu RECEPTOR (EFR), which recognize the bacterial MAMP epitopes flg22 and elf18, respectively (Gómez-Gómez and Boller 2000; Bauer et al. 2001; Kunze et al. 2004; Zipfel et al. 2006). FLS2 and EFR require co-receptors from the SERK family of leucine-rich repeat-receptor kinases, such as BAK1, to initiate immune signaling (Chinchilla et al. 2007; Tang et al. 2017). FLS2 is conserved across many plant species, whereas EFR is restricted to the Brassicaceae family (Tang et al. 2017). These PRRs have been instrumental in dissecting plant immunity, with assays frequently performed in both seedlings and rosettes. However, immune responsiveness increases during development across multiple species (Zhao et al. 2009; Wang et al. 2016). Additional outputs, such as flg22-induced callose deposition, also exhibit developmental regulation even when receptor abundance remains unchanged (Hu et al. 2023). Collectively, these findings indicate that developmental stage can substantially affect the magnitude and reliability of PRR-mediated immune responses.

Despite the widespread use of PRR-triggered immune assays, the extent to which assay type alters sensitivity across common immune readouts between receptor kinases has not been systematically evaluated. Common experimental systems include 10 to 14-day-old seedlings and 4 to 6-week-old rosettes. Seedlings are typically grown under long-day conditions on sterile Murashige Skoog (MS) medium, whereas rosette plants are grown under short-day conditions in soil under non-sterile conditions. These differences can substantially influence immune responses. Here, we systematically compare common immune outputs from FLS2 and EFR across *Arabidopsis* seedlings and rosettes under different experimental conditions. Our investigations reveal that assay conditions strongly influence the sensitivity and strength of immune responses. Careful consideration of how experimental conditions influence different immune assays is essential to determine the most appropriate system and MAMP concentration for obtaining robust and biologically meaningful results.

## Results

### PRR immune assays are widely represented in the literature

To evaluate the prevalence of primary research investigating PRR-mediated immunity, we performed Web of Science searches excluding review articles. Using the search terms “plant” AND “immunity” AND “receptor” AND (“MAMP” OR “PAMP”), we observed a strong increase in publications focused on PRR-mediated immunity since 1970, with publication rates peaking between 2020 and 2025 (Fig. 1A). Next, we evaluated the use of different plant species by performing additional searches incorporating species names. This analysis revealed that *Arabidopsis thaliana* is the most frequently studied, with approximately 2,889 publications, followed by *Nicotiana benthamiana* (1,020), *Oryza sativa* (920), *Solanum lycopersicum* (613), *Triticum aestivum* (376), and *Nicotiana tabacum* (348) (Fig. 1B). Together, these results emphasize both the widespread interest in PRR-mediated immunity and the dominant use of *Arabidopsis* as a model system for immune signaling studies.

**Figure 1.**
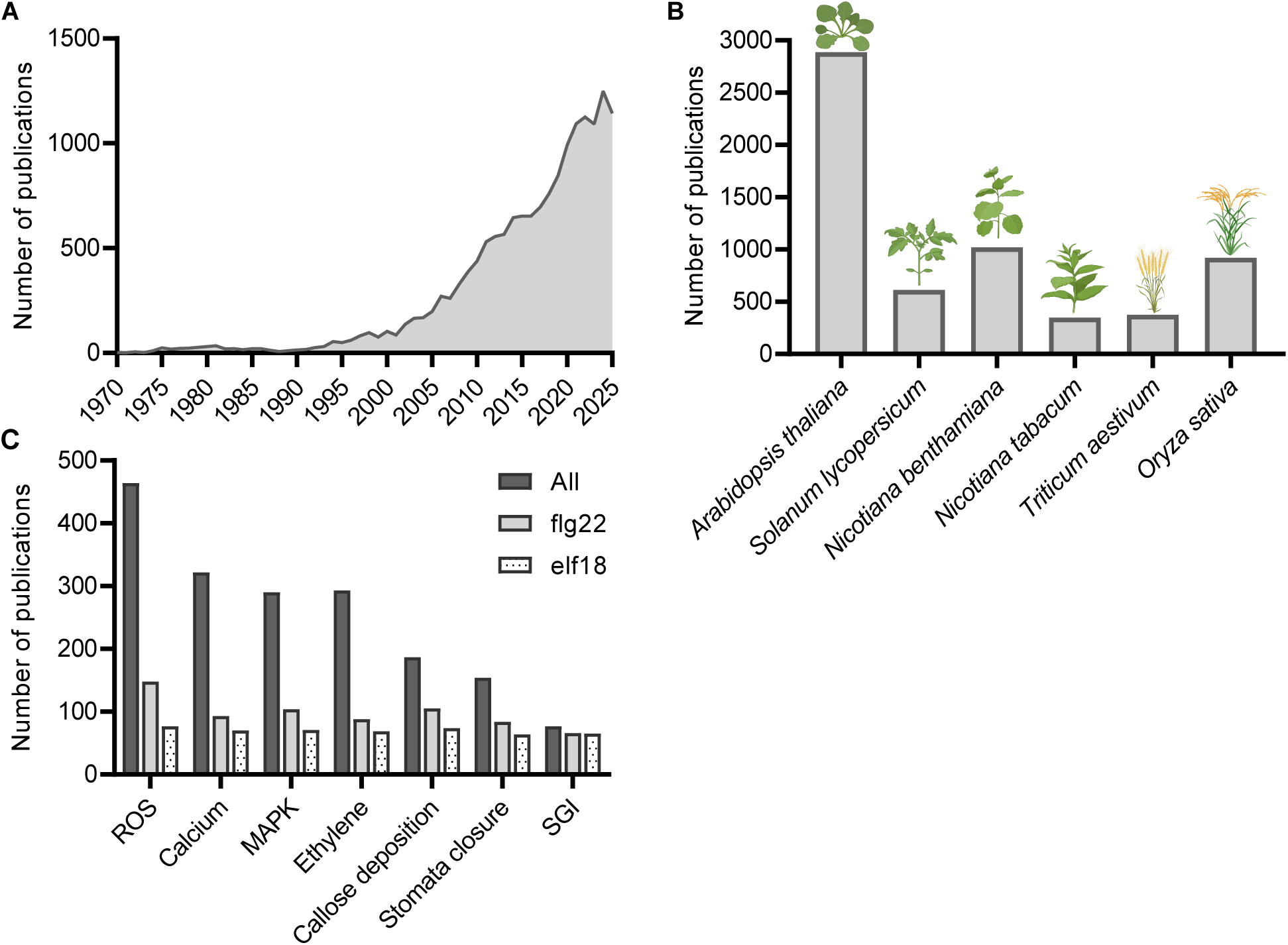
Publications on pattern-triggered immunity and immune assay usage. Charts represent the number of research articles that include the terms “plant” AND “ immunity” AND “receptor” along with the terms “PAMP” OR “MAMP” based on Web of Science queries. **A)** Historical overview of publications on pattern-triggered immunity since 1970 using the search terms described above. **B)** Number of publications by plant species. The bar chart represents the number of research articles that include the base search terms and specific plant species. **C)** Number of publications using *Arabidopsis thaliana* from (B) that report different immune assays. The bar chart represents the number of hits for each assay (All) or assays that specifically mention the epitopes flg22 or elf18. Data were obtained from Web of Science (https://www.webofscience.com/wos/woscc/smart-search). Plant images created in BioRender. Moreno perez, A. (2026), https://BioRender.com/tacyh90.

Next, we sought to determine which immune assays are most frequently used to evaluate MAMP perception in *Arabidopsis*. Web of Science searches incorporating *Arabidopsis thaliana* with assay-specific terms revealed that ROS production is the most frequently reported readout, followed by cytosolic calcium influx and MAPK activation (Fig. 1C). When searches were restricted to studies specifying individual MAMPs, flg22-associated publications were consistently more numerous than those using elf18 (Fig. 1C). Together, these results highlight the widespread reliance on a small set of standardized immune assays.

### Rosettes exhibit enhanced ROS sensitivity compared to seedlings

ROS production was the most frequently reported immune readout in *Arabidopsis* PRR studies (Fig. 1C). Because ROS can be measured rapidly and inexpensively using a luminol-based assay, it is widely used to evaluate MAMP immunogenicity (Boller and Felix 2009; Yu et al. 2017; Saeed and Trujillo 2022). ROS assays are frequently performed using either ∼14-day-old seedlings grown on MS medium or 4–6-week-old soil-grown rosettes (Mersmann et al. 2010; Colaianni et al. 2021; Stevens et al. 2024). For ROS measurements, we used detached true leaves from *Arabidopsis* seedlings and leaf discs from rosettes, as previous studies demonstrated that ROS responses differ between intact seedlings and excised leaf tissue (Mersmann et al. 2010).

We tested ROS responses to elf18 and flg22 in 14-day-old seedlings grown on MS medium and in 4-5-week-old soil-grown rosettes across a tenfold serial dilution series (100 nM to 0.01 nM; Fig. 2). The highest concentration we tested (100 nM) is commonly used in PRR immune assays; however, reported MAMP concentrations in the literature range widely, from 1 µM to 1 nM (Zipfel et al. 2006; Bredow et al. 2019; Dong et al. 2023). Seedlings treated with elf18 exhibited similar ROS kinetics and peak amplitudes at concentrations between 100 nM and 1 nM (Fig. 2, A and B). However, at 0.1 nM elf18, ROS production decreased by 83% relative to the 100 nM treatment (Fig. 2C). In contrast, rosettes maintained robust ROS production across a broader concentration range (100 nM to 0.1 nM; Fig. 2, A and B), with a 66% reduction only observed at 0.01 nM compared with 100 nM (Fig. 2C). Rosettes produced a strong ROS burst at 0.1 nM elf18, a concentration at which seedling responses were not significantly different from water controls. Thus, rosettes displayed approximately tenfold greater sensitivity to elf18 than seedlings.

**Figure 2.**
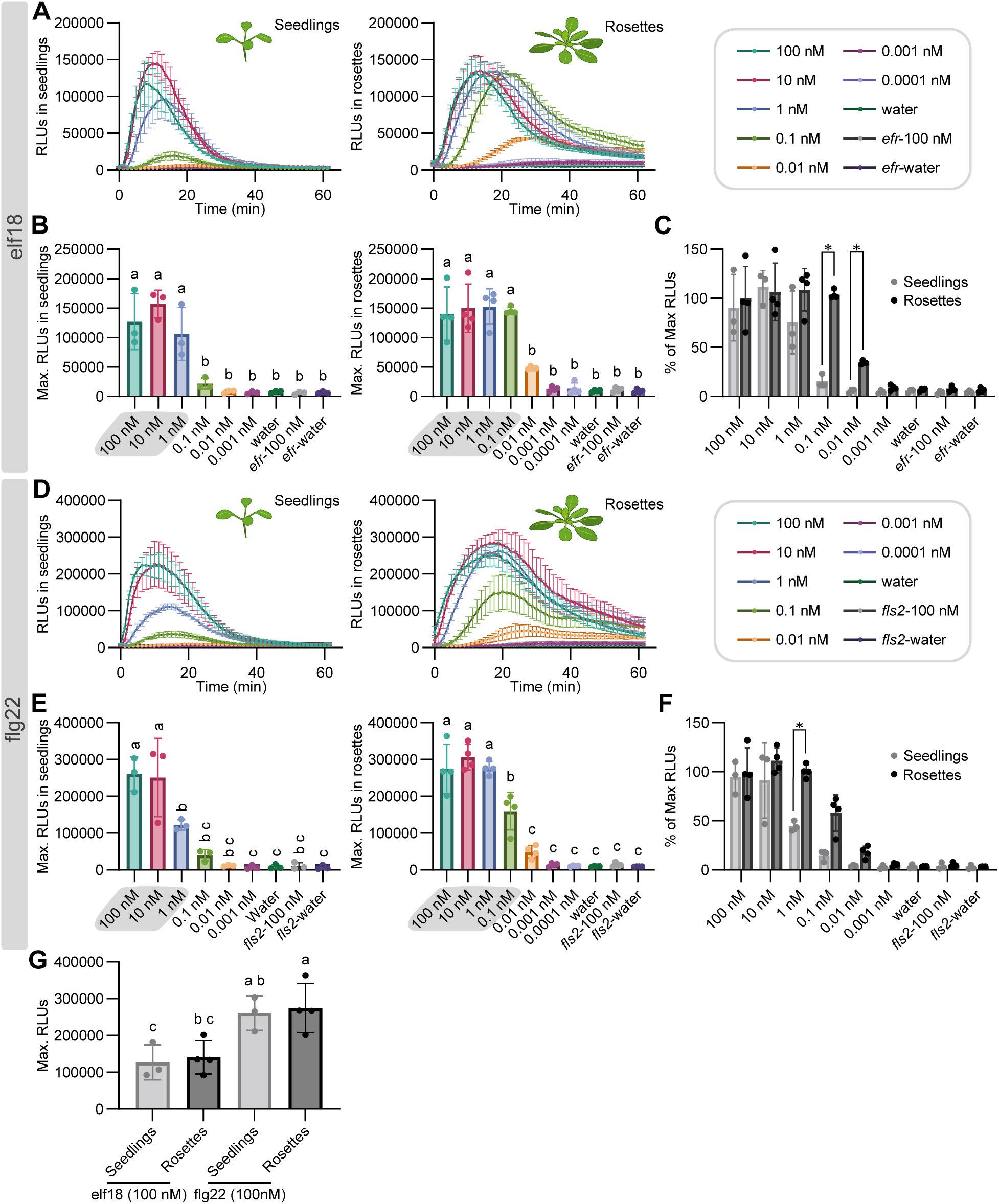
Plant age influences reactive oxygen species (ROS) sensitivity to elf18 and flg22. ROS production was analyzed in *Arabidopsis thaliana* Col-0 seedlings (14-day-old) and adult plants (4-5-week-old) treated with elf18 or flg22 at the indicated concentrations. As negative controls, the corresponding *efr-1* and *fls2-1* receptor mutants were included. Plant numbers and statistical analyses are identical for each peptide treatment. **A)** ROS burst kinetics in response to elf18. Curves represent mean luminescence (relative light units; RLU) ± SEM from 3 plates for seedlings (n = 24 seedlings) and 4 plates for adult plants (n = 8 plants). **B)** Quantification of maximum ROS production, using plants treated as described in (A). For seedlings, each data point represents the average max RLU from one plate containing eight seedlings (n = 24 seedlings). For adult plants, each data point represents the max average RLU from one plate containing four leaf discs from two plants (n = 8 plants). Error bars indicate SD. Statistical significances were determined by one-way ANOVA followed by Tukey’s HSD test (α = 0.05). **C)** ROS production expressed as a percentage of the maximal response observed at 100 nM elf18 in adult plants. Statistical significance was determined using multiple unpaired t-tests with Holm–Šídák correction for multiple comparisons applied to the raw data. **D)** ROS burst kinetics in response to flg22. **E)** Quantification of maximum ROS production, using plants treated as described in (D). **F)** ROS production expressed as a percentage of the maximal response observed at 100 nM flg22 in adult plants. **G)** Comparison of maximum ROS production at 100 nM elicitor concentration between elf18 and flg22 in seedlings and adult plants. Statistical significance was assessed using one-way ANOVA followed by Tukey’s HSD test (α = 0.05). Gray shading indicates concentrations that are significantly different from the water control.

Responses to flg22 showed a similar overall pattern. Seedlings exhibited ROS production between 100 nM and 1 nM flg22 (Fig. 2, D and E) but were less sensitive to flg22 than to elf18. At 1 nM flg22, seedling ROS production was 53% lower than at 100 nM (Fig. 2F). Rosettes responded to flg22 concentrations between 100 nM and 0.1 nM, although the 0.1 nM response was 42% lower than the 100 nM treatment (Fig. 2, D–F). Rosettes produced a robust ROS burst at 0.1 nM flg22, whereas seedling responses at this concentration were not statistically different from water controls. Therefore, rosettes displayed approximately tenfold greater sensitivity to flg22 than seedlings.

Both seedlings and rosettes responded strongly to elf18 and flg22 at 100 nM and 10 nM, indicating that high MAMP concentrations are sufficient to saturate ROS responses regardless of experimental conditions (Fig. 2, B and E). Across both experimental conditions, flg22 consistently triggered a stronger ROS burst than elf18 at 100 nM (Fig. 2G). However, elf18 triggered saturated ROS responses at ∼10-fold lower concentrations than flg22 in both seedlings and rosettes (Fig. 2, B and E). Together, these results demonstrate that rosettes exhibit markedly greater sensitivity to MAMP-induced ROS production than seedlings, and that elf18 elicits saturated responses at lower concentrations than flg22 in both experimental systems.

### Calcium reporter output is stronger in seedlings than rosettes

Cytosolic calcium influx is a rapid and quantitative marker of PRR activation (Yu et al. 2017). Unlike ROS measurements, calcium responses cannot be assessed directly in wild-type plants and require a transgenic reporter system. We used the established *Arabidopsis* aequorin line (Aeq^cyt^/pMAQ2) in the Col-0 background, which expresses cytosolic apoaequorin from the jellyfish *Aequorea victoria* (Knight et al. 1991). Upon Ca²⁺ binding, aequorin produces luminescence that can be quantified using plate-based luminometry (Ranf et al. 2012; Tanaka et al. 2013). Calcium reporter assays are commonly performed using ∼10-day-old seedlings grown on MS medium, whereas rosette assays are less frequently reported despite their widespread use for other immune outputs (Ranf et al. 2012; Maintz et al. 2014).

We tested calcium responses to elf18 and flg22 in 10-day-old seedlings grown on MS medium and in 4–5-week-old soil-grown rosettes across a range of MAMP concentrations (Fig. 3). The highest concentration we tested (100 nM) is frequently used in aequorin-based calcium assays; however, common MAMP concentrations range widely from 1 µM to 100 nM (Ranf et al. 2012; Monaghan et al. 2015). We applied tenfold serial dilutions to evaluate assay sensitivity. For elf18, seedlings displayed similar calcium kinetics and peak amplitudes at 100 nM and 10 nM (Fig. 3, A and B). Notably, 10 nM elf18 represented the lowest concentration that induced a detectable calcium response in both seedlings and rosettes, but the signal in seedlings was approximately 4.5-fold higher than in rosettes (Fig. 3, A and B). Across concentrations, calcium responses in rosettes were consistently reduced, with an approximately 78% lower signal than seedlings at 100 nM elf18 (Fig. 3C).

**Figure 3.**
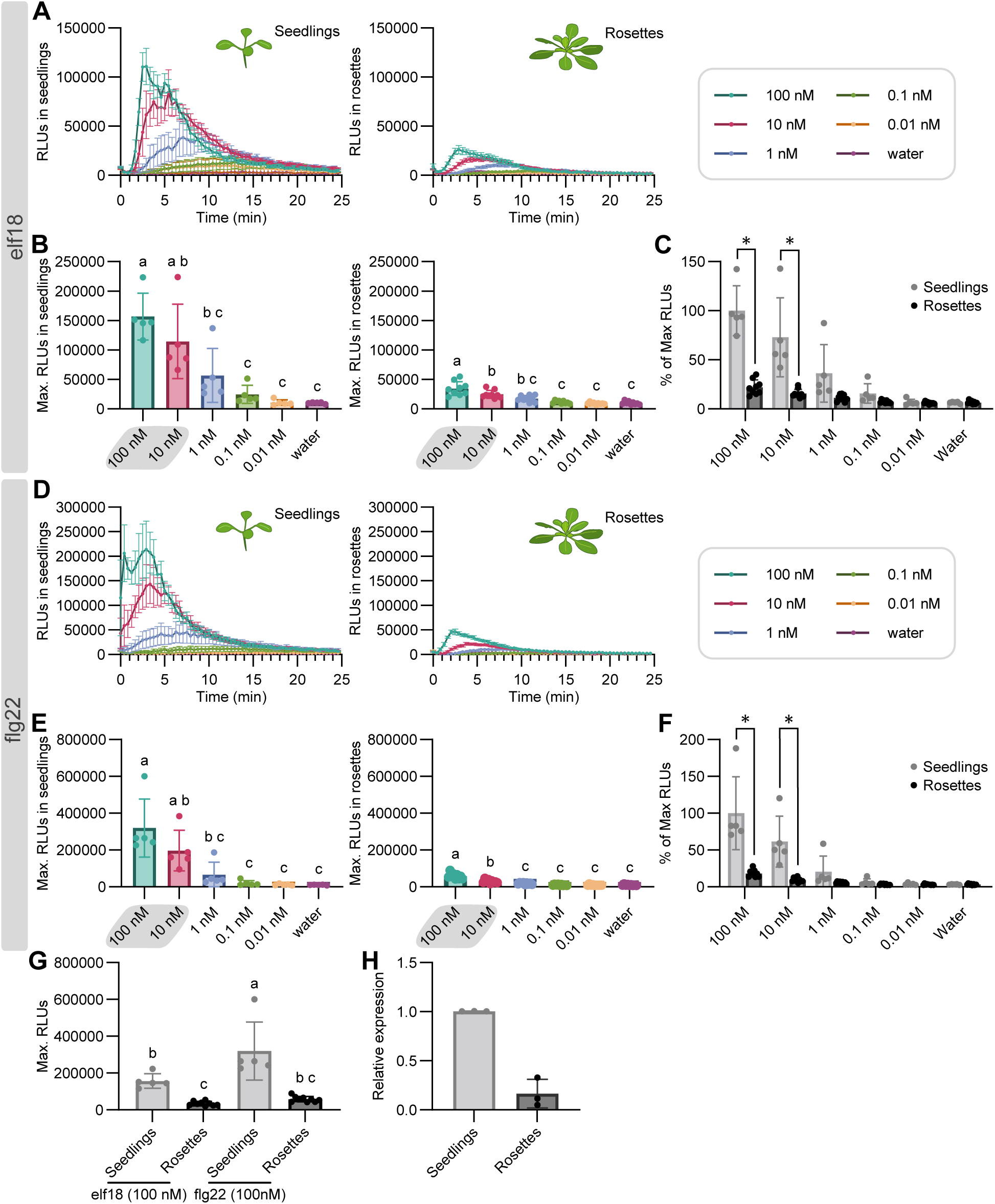
Plant age influences cytosolic calcium signaling and reporter expression. Cytosolic calcium responses were analyzed in *Arabidopsis thaliana* Col-0 plants expressing cytosolic p35S-apoaequorin (Aeq^cyt^/pMAQ2). Seedlings (10-day-old) and adult plants (4 to 5-week-old) were treated with elf18 or flg22 at the indicated concentrations. Plant numbers and statistical analyses are identical for each peptide treatment. **A)** Calcium burst kinetics in response to elf18. Curves represent mean luminescence (relative light units; RLU) ± SEM, from 5 plates for seedlings (n = 40 seedlings) and 9 plates for adult plants (n = 18 plants). **B)** Quantification of maximum calcium influx using plants treated as described in (A). For seedlings, each data point corresponds to the average max RLU from one plate containing eight seedlings (n = 40 seedlings). For adult plants, each data point represents the max average RLU from one plate containing four leaf discs from two plants (n = 18 plants). Error bars indicate SD. Statistical significance was assessed using one-way ANOVA followed by Tukey’s HSD test (α = 0.05). **C)** Calcium responses expressed as a percentage of the maximal response observed at 100 nM elf18 in seedlings. Statistical significance was determined using multiple unpaired t-tests with Holm–Šídák correction for multiple comparisons applied to the raw data. **D)** Calcium burst kinetics in response to flg22. **E)** Quantification of maximum calcium influx. **F)** Calcium responses expressed as a percentage of the maximal response observed at 100 nM flg22 in seedlings. **G)** Comparison of maximum calcium influx at 100 nM elicitor concentration between elf18 and flg22 in seedlings and adult plants. Statistical significance was assessed using one-way ANOVA followed by Tukey’s HSD test (α = 0.05). **H)** Transcript expression of apoaequorin in seedlings and adult plants of *Arabidopsis* Aeq^cyt^/pMAQ2 Col-0 background. cDNA samples from 10-day-old seedlings or 4-5-week-old plants were subjected to qPCR. Values represent relative expression compared to seedlings, means ± SD (n = 3). Data were analyzed using the ΔΔCt method and normalized against *Arabidopsis ELONGATION FACTOR 1α* (EF1α, AT5G60390). Significant differences were analyzed by Welch’s t-test (P < 0.05). Gray shading indicates concentrations that are significantly different from the water control.

Flg22 responses followed a similar trend. The minimal flg22 concentration that induced a significant calcium increase was 10 nM in seedlings and rosettes (Fig. 3, D and E). However, at 100 nM flg22, calcium reporter output was substantially higher in seedlings than in rosettes (Fig. 3F). As observed for ROS, flg22 triggered stronger calcium responses than elf18 at 100 nM in both experimental conditions (Fig. 3G). Unlike ROS assays, however, calcium reporter outputs did not reach comparable maximal levels in seedlings and rosettes, suggesting that reporter expression may differ between experimental conditions. To test this possibility, we quantified apoaequorin transcript abundance by qPCR. Reporter transcript levels were approximately 84% lower in rosettes than in seedlings (Fig. 3H). Together, these results indicate that calcium reporter output is substantially higher in seedlings than in rosettes, but this difference is largely attributable to reduced reporter expression in rosette tissue rather than intrinsic differences in calcium signaling sensitivity.

### Rosettes exhibit stronger MAPK activation than seedlings

We next analyzed MAPK activation, which occurs 5–15 minutes after elicitation by diverse immunogenic MAMPs (Yu et al. 2017). MAPK activation can be monitored by immunoblotting using anti-phospho ERK antibodies. These antibodies react with *Arabidopsis* MAPKs MPK3, MPK4, and MPK6, and detect homologous MAPKs in other plant species (Willmann et al. 2014). We compared elf18- and flg22-induced MAPK activation in 14-day-old seedlings grown in liquid MS medium and in 5–6-week-old soil-grown rosettes across a range of MAMP concentrations (Fig. 4). Although 1 µM is most commonly used for MAPK activation assays in adult tissue, reported MAMP concentrations typically range from 1 µM to 10 nM (Ranf et al. 2014; George et al. 2023). We therefore applied tenfold serial dilutions from 1 µM to 0.1 nM to assess assay sensitivity. MAPK activation was assessed 15 min after treatment, corresponding to the peak phospho-MAPK signal.

**Figure 4.**
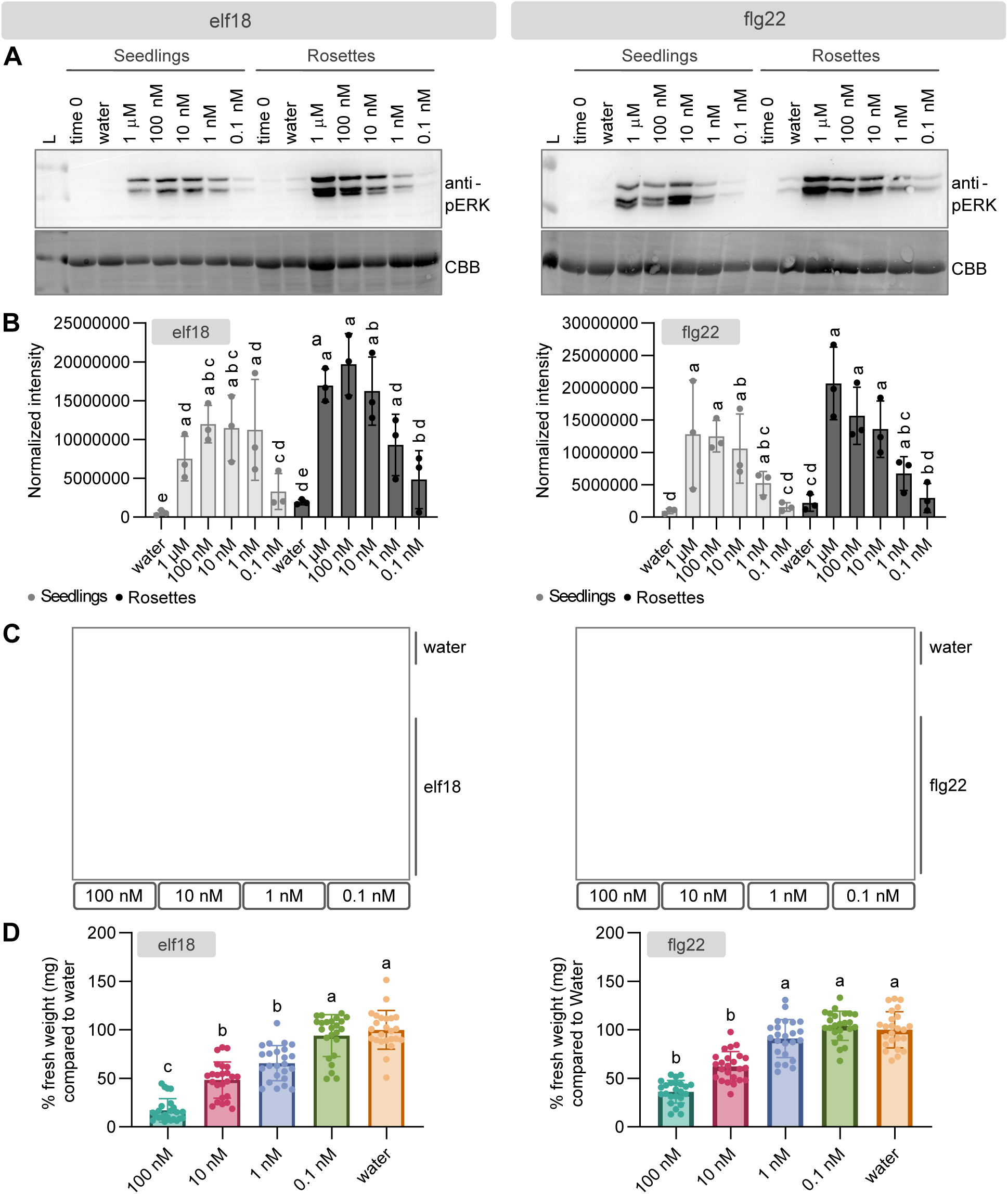
Plant age influences MAPK activation and seedling growth inhibition is more sensitive to elf18 than flg22. MAPK activation was analyzed in *Arabidopsis thaliana* Col-0 seedlings (14-day-old) and adult plants (4 to 5-week-old) treated with bacterial MAMP peptides at the indicated concentrations. Plant numbers and statistical analyses are identical for each peptide treatment. **A)** Induction of MAPKs by different concentrations of elf18 in Col-0 at 0 and 15 min after incubation in response to elf18 (left) and flg22 (right). CBB = protein loading. All experiments were repeated at least three times with similar results. **B)** Quantification of band intensity of MAPK activation induced in response to elf18 and flg22 on A). Each data point corresponds to the normalized band intensity from one independent blot ± SD (n = 3). The MAPK immunoblot were quantified in Image Lab Software (Bio-Rad), and samples were normalized to the intensity of the Coomassie blue staining. Statistical differences were determined by Lognormal ordinary one-way ANOVA followed by Tukey’s test (α = 0.05), letters indicates significant differences. **C)** *Arabidopsis* SGI in response to elf18 (left) or flg22 (right) at the indicated concentrations. **D)** Bar charts represent SGI from C) expressed as a percentage relative to the water-treated control, which was set to 100% (maximum growth). Each data point represents an individual plant. Eight plants were grown per plate for each condition, and data were pooled from three independent experiments (n = 24 seedlings). Statistical significance was assessed using Krustal-Wallis and Dunn’s multiple comparison test (P < 0.0001).

For elf18, MAPK activation was detectable at all concentrations seedlings, with signal intensity gradually decreasing at lower MAMP concentrations (Fig. 4, A and B). Across concentrations, rosettes consistently exhibited stronger elf18-induced MAPK activation than seedlings, indicating enhanced responsiveness in rosette experimental conditions (Fig. 4, A and B). flg22-induced MAPK activation was not detectable below 1 nM (Fig. 4, A and B). Direct comparisons between elf18- and flg22-induced MAPK signal intensity are not possible because samples were not run on the same membrane. Moreover, immunoblot quantification has a limited dynamic range that is influenced by antibody sensitivity and signal saturation and should therefore be interpreted as semi-quantitative (Aldridge et al. 2008; Taylor and Posch 2014). Together, these results show that rosettes exhibit stronger MAPK activation than seedlings, and that elf18 induces detectable responses at lower concentrations than flg22 in seedlings.

### Elf18 induces stronger seedling growth inhibition than flg22

To examine a later stage of the immune response, we analyzed SGI, a downstream output that can be measured 8-10 days after elicitation in *Arabidopsis* and reflects a reduction in growth associated with sustained immune activation (Boller and Felix 2009; Yu et al. 2017). SGI assays are typically performed using seedlings grown on MS medium supplemented with immunogenic MAMPs.

We tested the ability of elf18 and flg22 to induce SGI in 12-day old *Arabidopsis* seedlings that were treated with a range of MAMP concentrations for 8 days. Reported peptide concentrations for SGI assays range from 10 µM to 1 nM (Gómez-Gómez et al. 1999; Ranf et al. 2011; Bredow et al. 2019). The highest concentration tested (100 nM) is commonly used for SGI assays, and tenfold serial dilutions were applied to evaluate sensitivity, (Fig. 4, C and D). Elf18 induced significant SGI from 100 nM to 1 nM, whereas flg22 triggered significant inhibition only down to 10 nM. Thus, seedlings displayed approximately tenfold greater sensitivity to elf18 than to flg22. This contrasts with early immune outputs, where flg22 elicited stronger ROS and calcium responses.

## Discussion

Here, we provide a systematic comparison of multiple PRR-mediated immune outputs across seedling and rosette experimental systems and across a range of flg22 and elf18 concentrations. Our results demonstrate that immune output sensitivity is strongly influenced by assay type, developmental stage, and growth conditions. Calcium signaling, ROS production, and MAPK activation can be measured in both seedlings and rosettes, but their application differs widely across the literature. ROS assays are most often performed using rosette leaf tissue, whereas calcium signaling and MAPK activation are frequently assessed in seedlings using different growth substrates and photoperiods (Ranf et al. 2011, 2012; Thor and Peiter 2014; Ma et al. 2021; Dong et al. 2023; Stevens et al. 2024). Studies combine data obtained from rosettes and seedlings within the same work, performing different assays at different developmental stages.

Our results show that immune responses can be saturated at commonly used MAMP concentrations, yet sensitivity differences become apparent at lower doses. In the literature, a wide range of MAMP concentrations is reported (10 nM to 1 µM), with only a limited number of studies employing concentrations below 10 nM (Bauer et al. 2001; Zipfel et al. 2006; Bredow et al. 2019). During natural infection, pathogen distribution is dynamic and heterogeneous, and MAMP expression, such as flagellin, can be downregulated over time and across tissues, generating localized differences in MAMP abundance (Zhu et al. 2023; López-Pagán et al. 2025). Because MAMP variants can display reduced immunogenic activity (Colaianni et al. 2021; Parys et al. 2021; Stevens et al. 2024), minimal effective concentrations should be empirically determined for each peptide. Based on our results, we suggest 10–100 nM as a practical working range for flg22 and elf18 assays and recommend avoiding micromolar peptide concentrations.

Age-related resistance, defined as the progressive increase in disease resistance with plant development, has been observed across multiple species (Hu and Yang 2019; Hu et al. 2023; Li et al. 2024). Developmental stage impacts PRR-mediated immunity (Hu and Yang 2019; Li et al. 2024). For example, expression of the csp22 receptor *CORE* (Cold Shock Protein Receptor) increases with age in *Nicotiana benthamiana*, correlating with enhanced csp22-induced defenses (Wang et al. 2016). Similarly, the rice receptor *Xa21* confers weak resistance in juvenile plants but strong immunity at later developmental stages due to age-dependent accumulation of *Xa21* transcripts (Zhao et al. 2009). In *Arabidopsis*, the microRNA miR172 promotes increased *FLS2* expression and enhanced downstream PRR-immune responses during microbiota early seedling development, particularly between 3- and 6-day-old plants (Zou et al. 2018). Consistent with age-dependent changes in immunity, we observed stronger MAPK activation and increased ROS sensitivity in rosettes compared to seedlings. However, developmental stage is not the only variable differentiating seedling and rosette experimental systems. These also differ in photoperiod, microbiota, and growth substrate. Long day conditions positively influence the timing of age-related resistance in *Arabidopsis* (Rusterucci et al. 2005). In addition, axenically grown plants exhibit defects in immune competence and reduced age-dependent resistance, suggesting that microbiota composition contribute to developmental regulation of immunity (Paasch et al. 2023).

In contrast to ROS production and MAPK activation, calcium responses were weaker in rosettes than in seedlings. Calcium influx measurements rely on the transgenic Aeq^cyt^/pMAQ2 reporter line (Knight et al. 1991, 1996; Ranf et al. 2012). Apoaequorin transcript abundance was approximately 84% lower in rosettes than in seedlings, consistent with the reduced calcium signals observed in adult tissue. The reporter is driven by the viral CaMV 35S promoter, which exhibits developmental, tissue-specific, and insertion site–dependent variability, with higher activity often observed in younger tissues than in mature leaves across multiple species (Pret’ová et al. 2001; van Leeuwen et al. 2001; Amack and Antunes 2020; Kiselev et al. 2021). Most calcium signaling experiments using the Aeq^cyt^/pMAQ2 line have been performed in seedlings or in protoplasts derived from rosette tissue (Ranf et al. 2012; Maintz et al. 2014). Calcium signals have also been reported in rosette leaf discs with higher MAMP and coelenterazine concentrations (Jeworutzki et al. 2010; Monaghan et al. 2015). Together, these findings highlight that reporter expression is not uniform over time and calcium reporter outputs should be interpreted cautiously when comparing signaling across plant developmental stages and experimental conditions.

The amplitude and strength of immune responses can also vary during perception of distinct MAMPs under identical experimental conditions (Ranf et al. 2011; Wan et al. 2019; Robinson et al. 2025). For example, perception of nlp20 by the receptor protein RLP23 and flg22 by the receptor kinase FLS2 trigger similar immune responses but differ in the kinetics and magnitude of Ca²⁺ influx, ethylene production, and callose deposition (Wan et al. 2019). Consistent with previous studies (Zipfel et al. 2006; Ranf et al. 2011), we found that flg22 elicits stronger early immune responses, including ROS production and calcium influx, than elf18. However, a tenfold lower concentration of elf18 is sufficient to reach a saturated ROS response. In contrast, elf18 more effectively induces SGI, indicating that stronger early signaling does not necessarily predict downstream growth inhibition. This distinction may reflects differences in growth regulation, as flg22 preferentially restricts root growth whereas elf18 has a stronger effect on shoot growth, resulting in a greater reduction in whole-seedling biomass (Ranf et al. 2011). Tissue specificity also contributes to differences in response amplitude among MAMPs. For example, isolated *Arabidopsis* roots produce ROS in response to flg22 but not elf18 (Wyrsch et al. 2015), *FLS2* expression is regulated by damage in the root (Zhou et al. 2020), and seedling root calcium influx is triggered by the chitin octamer and Pep1 phytocytokine but not by flg22 or elf18 (Ranf et al. 2011).

We provide a systematic comparison of commonly used immune assays in *Arabidopsis* seedlings and rosettes in response to flg22 and elf18. Our results demonstrate that canonical PRR-mediated immune outputs are strongly influenced by assay type, growth conditions, and reporter systems, and should not be interpreted as interchangeable measures of immune activation. Based on these findings, we recommend evaluating MAMP sensitivity when comparing tissues or developmental stages, integrating multiple immune outputs, and using the lowest MAMP concentration that yields robust and reproducible responses. We anticipate that this work will serve as a practical resource for improving the experimental design and interpretation of plant immune assays.

## Methods

### Literature searches

Publication data were obtained from Web of Science (Core Collection; https://www.webofscience.com) using the Advanced Search function. For all searches, the document type was restricted to “Article,” and terms were queried within the “Topic” field. The base search included the terms “plant” AND “immunity” AND “receptor” combined with “PAMP” OR “MAMP.” For the historical overview, the number of publications per year was searched on January 30, 2026.

To determine publication numbers by plant species, the base search was combined with the names of individual species, including *Arabidopsis thaliana*, *Nicotiana benthamiana*, *Nicotiana tabacum*, *Oryza sativa*, *Solanum lycopersicum*, and *Triticum aestivum*. To assess the prevalence of specific immune assays, the base search and *Arabidopsis thaliana* were combined with assay-related terms, including “reactive oxygen species,” “calcium,” “callose deposition,” “ethylene,” “mitogen-activated protein kinase,” “seedling growth inhibition,” and “stomatal closure.” All search results were saved as HTML files for documentation.

### Plant Material and Growth Conditions

For seedling experiments, *Arabidopsis thaliana* ecotype Columbia (Col-0) and their transgenic lines *efr-1* (SALK_044334) (Zipfel et al. 2006), *fls2-T* (SALK_062054) (Ranf et al. 2012), and Aeq^cyt^/pMAQ2 (Knight et al. 1991, 1996; Ranf et al. 2012) were surface-sterilized with bleach and stratified at 4°C for 2–3 days. Seeds were then germinated on one-half-strength Murashige and Skoog medium (½ MS) (Research Products International, Mount Prospect, IL, USA) plates containing 1% sucrose and 0.8% plant agar. Seedlings were grown in a growth chamber under a 16-h light/8-h dark photoperiod at 22°C, and ∼100 μmol m⁻² s⁻¹ light intensity). The ages chosen for seedling assays were based on those commonly used in previous studies, and the specific age used for each assay is indicated below.

For rosettes, *A. thaliana* Col-0 and the transgenic lines *efr-1* and *fls2-T* were grown in Sunshine Mix #1 soil (Sun Gro Horticulture, Agawam, MA, USA) under controlled conditions (10-h light/14-h dark cycle, 23°C, ∼100 μmol m⁻² s⁻¹ light intensity, and 70% relative humidity). To ensure proper germination, Aeq^cyt^/pMAQ2 seedlings were germinated on MS medium and transplanted to soil four days after germination. As with seedlings, plant ages for rosette assays were selected based on previous studies, and the exact age is indicated in each experiment below.

### MAMP peptides

The MAMP peptides elf18 (acyl-SKEKFERTKPHVNVGTIG) were ordered from (Genscript, Piscataway, New Jersey, USA) and flg22 (QRLSTGSRINSAKDDAAGLQIA) were ordered from (Shanghai Apeptide CO., LTD, Shanghai, China) with purities of 97.6% and 95.5%, respectively. Peptides were dissolved in sterile water prior to use.

### Measurement of Reactive Oxygen Species (ROS)

For ROS assays, *A. thaliana* Col-0, *efr-1*, and *fls2-T* seeds were germinated on ½ MS plates for four days. Seedlings were then transferred to a Fisherbrand™ 96-well tissue culture plate (FisherScientific, Cat. No. FB012931) containing 150 μL of liquid ½ MS medium. Plates were sealed with 3M Micropore Surgical Tape (Cat. No. 1530-1) and returned to the growth chamber. After ten days of growth, the true leaves of 14-day-old seedlings were used for assays (Mersmann et al. 2010).

For experiments with rosettes, leaf punches (3 mm diameter) were taken from equivalent leaves of four- to five-week-old plants using a biopsy punch (Stevens et al. 2024). True leaves of seedlings or leaf disks from rosettes were floated abaxial side down in 190 μL of sterile water in white 96-well microtiter plates (Costar^®^, ref 3912) and incubated for 18–24 h in dark.

To elicit ROS production, water was replaced with 100 μL of assay solution containing sterile water, 20 μM L-012 (Wako Chemicals, Cat. No. 143556-24-5), 20 μg/mL horseradish peroxidase (HRP; Thermo Fisher, Cat. No. 31490), and the indicated concentration of MAMP peptide. Assay measurements were taken every 1.25 minutes for 61.25 minutes. Luminescence was recorded over time using a plate reader (BioTek Synergy H1, Agilent, Santa Clara, California, USA).

### Aequorin Luminescence Measurements (Cytosolic Calcium Assay)

For cytosolic calcium measurements, *A. thaliana* Col-0 plants expressing p35S-apoaequorin (pMAQ2) in the cytosol (Aeq^cyt^) were germinated on ½ MS plates for four days. Seedlings were then transferred to a 96-well plate (FisherScientific, Cat. No. FB012931) containing 150 μL of liquid ½ MS medium. Plates were sealed with 3M Micropore Surgical Tape (Cat. No. 1530-1) and returned to the growth chamber. Ten-day-old seedlings were used for seedling-based assays (Ranf et al. 2012). For rosette-stage experiments, seeds were germinated on ½ MS medium for 7 days and then transferred to soil; 4–5-week-old rosette plants were used for all experiments (Monaghan et al. 2015). True leaves from 10-day-old seedlings or 3-mm-diameter leaf discs from rosettes were incubated overnight (18–24 h) in 100 μL of 5 μM coelenterazine (CTZ; Cat. No. CZ2.5, Gold biotechnology, St. Louis, MO, USA) in the dark. To measure calcium influx, CTZ solution was replaced with 100 μL of peptide solution at the indicated concentrations prepared in sterile water. Luminescence was recorded immediately after elicitation. Assay measurements were taken every 25 seconds for 25 minutes. Luminescence was recorded over time using a plate reader (BioTek Synergy H1, Agilent, Santa Clara, California, USA).

### MAPK Induction Assay

For MAPK activation assays, *A. thaliana* Col-0, *efr*, and *fls2* seeds were germinated on ½ MS plates for five days, then transferred to 48-well plates (Costar^®^, ref 3548) containing 600 μL of liquid ½ MS medium. Plates were sealed with 3M Micropore Surgical Tape and returned to the growth chamber. After nine days, 14-day-old seedlings were treated with sterile water or the indicated concentration of elf18 or flg22 for 0 or 15 min before harvesting. Three seedlings were pooled per treatment.

For rosettes, 7-mm leaf punches were collected from equivalent leaves of five- to six-week-old plants and transferred to 24-well plates (Costar^®^, ref 3526) containing 1 mL of sterile water. Each well contained three leaf discs collected from three biological replicates. Leaf discs were incubated overnight in sterile water and then treated with the indicated concentration of elf18 or flg22 for 0 or 15 min before harvesting.

Seedlings and leaf discs were frozen in liquid nitrogen, ground to a fine powder, and resuspended in 200 μL of extraction buffer (50 mM HEPES, pH 7.5; 50 mM NaCl; 10 mM EDTA; 0.2% Triton X-100; 1× Pierce protease inhibitor mini tablet [Thermo Scientific, Cat. No. A32955]; 1× Pierce phosphatase inhibitor mini tablet [Thermo Scientific, Cat. No. A32957]). Total proteins were isolated by centrifugation at 21,000 × *g* for 10 min at 4°C, and Laemmli buffer (without bromophenol blue) was added for SDS–PAGE.

Protein concentrations were determined using the Pierce 660 nm Protein Assay (Thermo Scientific, Cat. No. 22660) with Ionic Detergent Compatibility Reagent (Thermo Scientific, Cat. No. 22663). Equal amounts of protein were separated by 12% SDS–PAGE, and phosphorylated MAPK3 and MAPK6 were detected by immunoblotting using p44/42 MAPK (pERK) antibodies (Cell Signaling Technology, Cat. No. 4370; 1:2000 dilution) and secondary rabbit antibodies (Bio-Rad, Cat. No. 1705046; 1:3000 dilution). Antibodies were diluted in 1× TBS-T containing 5% BSA. Antibodies were detected with enhanced chemiluminescence using the SuperSignal^TM^ West Pico PLUS Chemiluminescent Substrate (Thermo Scientific, Cat. No. 34578). Experiments were repeated at least three times with consistent results. MAPK immunoblots were quantified in Image Lab Software (Bio-Rad), band intensities were normalized to the Coomassie Blue staining signal (Taylor and Posch 2014).

### Seedling Growth Inhibition (SGI) Assay

Seedling growth inhibition assays were performed as described by (Stevens et al. 2024). *A. thaliana* Col-0 and *efr* seeds were germinated on ½ MS plates for four days, then transferred to 48-well plates containing 500 μL of liquid ½ MS medium supplemented with the indicated concentration of elf18 or flg22. Plates were sealed with 3M Micropore Surgical Tape and maintained in the growth chamber. After eight days, seedlings were gently blotted dry and weighed. Each treatment included eight seedlings per replicate. Mock controls (½ MS medium only) were included on each plate. Experiments were repeated three times with similar results.

### RNA Extraction and Quantitative PCR (qPCR)

Total RNA was extracted from 14-day-old seedlings or four- to five-week-old rosettes of *A. thaliana* Col-0 expressing Aeq^cyt^/pMAQ2 using TRIzol reagent (Fisher Scientific, Cat. No. 15596018) following the manufacturer’s protocol. RNA was treated with RQ1 RNase-Free DNase (Promega, Cat. No. M6101) to remove genomic DNA contamination. Three biological replicates were analyzed per treatment. Complementary DNA (cDNA) was synthesized using M-MLV Reverse Transcriptase (Promega, Cat. No. M170A) and Oligo(dT)15 Primer (Promega, Cat. No. C110A) following the manufacturer’s protocol. SsoFast EvaGreen Supermix (Bio-Rad, USA) was utilized for qPCR reaction following the manufacturer’s protocol on a CFX96 Real-Time PCR Detection System (Bio-Rad). Thermocycling steps were as follows: 95 °C for 30 s followed by 40 cycles alternating between 5 s at 95 °C and 5 s at 60 °C. The melt curve was calculated after the final cycle by alternating between 65 °C for 5 s and 95 °C for 5 s. Relative transcript abundance was determined using the ΔΔCt method (Livak and Schmittgen 2001) and normalized to *Arabidopsis ELONGATION FACTOR 1α* (*EF1α*; AT5G60390), which served as the reference gene. Because one PCR cycle represents an approximate twofold difference in template abundance, fold changes were calculated as 2^−ΔΔCt^ (Livak and Schmittgen 2001). For amplification of *EF1α*, forward primer 5’-CTGGATTCGAGGGAGACAACA-3’ and reverse primer 5’-GCACCGTTCCAATACCACCAA-3’ were used. For amplification of *APOAEQUORIN* GenBank accession M16103.1), forward primer 5’-CCAGACTTCGACAACCCAAA-3’ and reverse primer 5’-TTGTTCAGGTGTTGCTCCAA-3’ were used.

### Statistical analysis

All statistical analyses were performed using GraphPad Prism v10 (GraphPad Software, San Diego, CA, USA). For comparisons among more than two groups with normally distributed data, statistical significance was assessed by one-way ANOVA followed by Tukey’s honestly significant difference (HSD) post hoc test (α = 0.05). For percentage statistical significance was determined using multiple unpaired *t*-tests with Holm–Šídák correction for multiple comparisons applied to the raw data. Comparisons between two groups were performed using Welch’s *t* test (P < 0.05). Statistical differences in MAPK activation were determined by Lognormal ordinary one-way ANOVA followed by Tukey’s test (*α* = 0.05). Statistical differences in SGI data, that did not meet normality assumptions, was evaluated using the Kruskal–Wallis test followed by Dunn’s multiple comparison test. The specific statistical test applied in each case is indicated in the corresponding figure legend. All experiments were repeated at least three times with similar results.

## Data Availability

All raw data underlying each figure have been deposited in Zenodo: 10.5281/zenodo.19863133

## Acknowledgments

This research was supported by the NIH grant R35GM136402 to G.C. A.M.P. was supported by the Fundación Alfonso Martín Escudero Postdoctoral fellowship. We also would like to thank Marta Bjornson and all the members of Coaker lab for helpful discussions and input on the project. We thank T. Chilcote for excellent technical assistance and Shahid Siddique (UC Davis) for the usage of the Biotek plate reader.

## Author contributions

G.C. conceived and supervised the study. A.M.P. contributed to conceptualization and performed most of the experiments. G.C. and A.M.P. analyzed the data. H.S. assisted with MAPK and ROS assays and plant growth. A.M.P. and G.C. wrote the manuscript.

## Notes

https://doi.org/10.5281/zenodo.19863133

### Competing Interest Statement

The authors have declared no competing interest.

